# High-throughput 3D super-resolution imaging in deep tissue

**DOI:** 10.1101/2025.03.07.641978

**Authors:** Zhenhong Du, Jiajia Chen, Cancan Gao, Jiayu Li, Qin Ru, Yao Zheng, Wei Gong, Ke Si

**Affiliations:** Department of Psychiatry of the First Affiliated Hospital and School of Brain Science and Brain Medicine, State Key Laboratory of Extreme Photonics and Instrumentation, Zhejiang University School of Medicine, Hangzhou 310058, China; Nanhu Brain-computer Interface Institute, Hangzhou 311100, China; College of Optical Science and Engineering, Zhejiang University, Hang Zhou 310027, China; Liangzhu Laboratory, MOE Frontier Science Center for Brain Science and Brain-machine Integration, State Key Laboratory of Brain-machine Intelligence, Zhejiang University, 1369 West Wenyi Road, Hangzhou 311121, China; NHC and CAMS Key Laboratory of Medical Neurobiology, Zhejiang University, Hangzhou 310058, China; Lingang Laboratory, Shanghai, 200031, China

**Author notes:** These authors contributed equally to this work (Zhenhong Du, Jiajia Chen, Cancan Gao).

## Abstract

Fluorescence images obtained with optical microscopes intrinsically suffer from blur and noise, which can be partially reversed by the deconvolution process. However, the deconvolution process is ill-conditioned, leading to a trade-off between detail preservation and noise suppression. Here, we develop 3D-FUDIP to fully decouple the deconvolution process into two parts: deblurring and denoising, achieving an 8-fold improvement in spatial resolution. By adopting the Poisson model, which obeys the quantum nature of photons, our 3D-FUDIP can be successfully applied to various noise conditions, especially low-light conditions where the photon number is generally extremely small. The results show that our 3D-FUDIP improves the SNR by up to 6-fold with only a one-fifth photon budget. Besides, 3D-FUDIP boosts the spatial bandwidth product (SBP) by one order of magnitude, allowing more spine details to be resolved within a larger imaging volume. By synergizing deep learning with these advances, we propose 3D-FUDIPn to further improve the imaging resolution. We demonstrate 3D-FUDIP’s performance in various imaging systems, including confocal, two-photon, and light-sheet microscopes, showing compatibility and potential applications in biological science.

## Introduction

Fluorescence imaging techniques have greatly facilitated the visualization and analysis of structures and functions of biological samples^1, 2^. Although the performance of fluorescence microscopy has been continuously pushed forward, fluorescence images still suffer from serious deterioration introduced by blur and noise^3, 4^. To recover a high signal-to-noise ratio (SNR), various denoising methods have been proposed, including gradient prior-based methods^5-7^, self-similarity prior-based methods^8-10^, and camera correction-based methods^11, 12^ etc. The gradient prior-based methods, such as TV^13^ and Hessian^5^ denoising, effectively suppress noise and artifacts in structure illumination microscopy. The self-similarity prior-based methods, such as non-local means (NLM)^14^ and block matching with 4D filtering (BM4D)^9^ algorithms, demonstrate good denoising capability for certain volumetric data. The camera correction-based methods, such as the noise correction algorithm for sCMOS camera (NCS)^11^ and the automatic correction of sCMOS-related noise (ACsN)^12^, promote effective suppression of raw fluorescence images from sCMOS cameras by accurately estimating camera characteristics. As a balance, these denoising methods usually sacrifice image details due to the trade-off between noise suppression and detail preservation^15^.

To partially reverse image degradation, various deconvolution techniques have been developed. Richardson-Lucy (RL) deconvolution^16^ effectively improved the resolution and contrast of fluorescence images through maximum likelihood estimation, while it is susceptible to noise interference^17^. To restore the latent structural information submerged by noise, total variation (TV) regularization was incorporated into RL deconvolution to preserve details in 3D microscopy^18^. The utilization of prior information in fluorescence imaging brought evident resolution improvement^13^. However, this approach induced the staircase effect^6^ simultaneously, which deteriorated the image quality to some extent. More recently, sparse deconvolution combining Hessian and *L*_1_ norm regularization methods leveraged the sparsity and continuity priors of fluorescence images to overcome the resolution limits of super-resolution microscopes^17^. Besides, the burgeoning deep neural networks (DNNs) show unique advantages in various image restoration tasks due to their powerful feature extraction capability^19-25^. However, current DNNs mainly rely on a supervised manner to achieve effective end-to-end mapping. This requires a large amount of high-quality ground truth^25^, which is often difficult or even impractical to obtain in fluorescence imaging^26^. On the other hand, unsupervised learning reduces the reliance on paired data by utilizing the intrinsic structure and statistical characteristics of the input data^27-32^. However, this potentially reduces the accuracy of results and increases the training complexity^33^.

To break the trade-off between noise suppression and fine detail preservation in fluorescence imaging, we propose a 3D fully uncoupled deconvolution method with alternating iteration based on the Poisson model (3D-FUDIP). By using variable splitting, we can fully decouple the deconvolution process into deblurring and denoising parts. We then further optimize the image performance through alternating iterations. By adopting the Poisson model, which obeys the quantum nature of photons^34^, our 3D-FUDIP can be successfully applied to various noise conditions, especially low-light conditions where the photon number is generally extremely small. The results show that our 3D-FUDIP improves the SNR by up to 6-fold and spatial-bandwidth product (SBP) by 20-fold. With our 3D-FUDIP, more spine details can be resolved with larger imaging volume, to some extent circumventing the trade-off between resolution and fields of view (FOVs). Further, we design 3D-FUDIPn to push forward the limit of spatial resolution by collaborating 3D-FUDIP with deep learning. The enhancement in axial resolution has enabled more accurate morphology reconstruction and classification of dendritic spines. Finally, we extend our 3D-FUDIP to various imaging systems, including confocal microscopy, two-photon microscopy, and light-sheet microscopy, showing potential applications in biological science.

## Results

### Principles of the 3D-FUDIP

During the fluorescence imaging process, the finite numerical aperture (NA) of the objective lens, aberrations, and noise inevitably lead to image degradation^35^. Traditional denoising methods focus on enhancing the SNR of images, at the price of losing details and blurring edges. Deconvolution methods can partially solve the blurring issues but usually induce noise amplification and artifacts.

To break the trade-off between detail preservation and noise suppression, we establish a physical model to describe the fluorescence imaging process and devise a 3D fully uncoupled deconvolution method, termed 3D-FUDIP (Fig. 1a). We apply sparsity constraints in the tight wavelet domain as a regularization term to preserve edge details of fluorescence volumes in 3D-FUDIP. Then we introduce a data-fitting term that explicitly constructs the deblurring and denoising challenges. Consequently, the deconvolution process is fully decoupled into the deblurring and denoising sub-problems through splitting variables and alternating iterations.

**Fig. 1.**
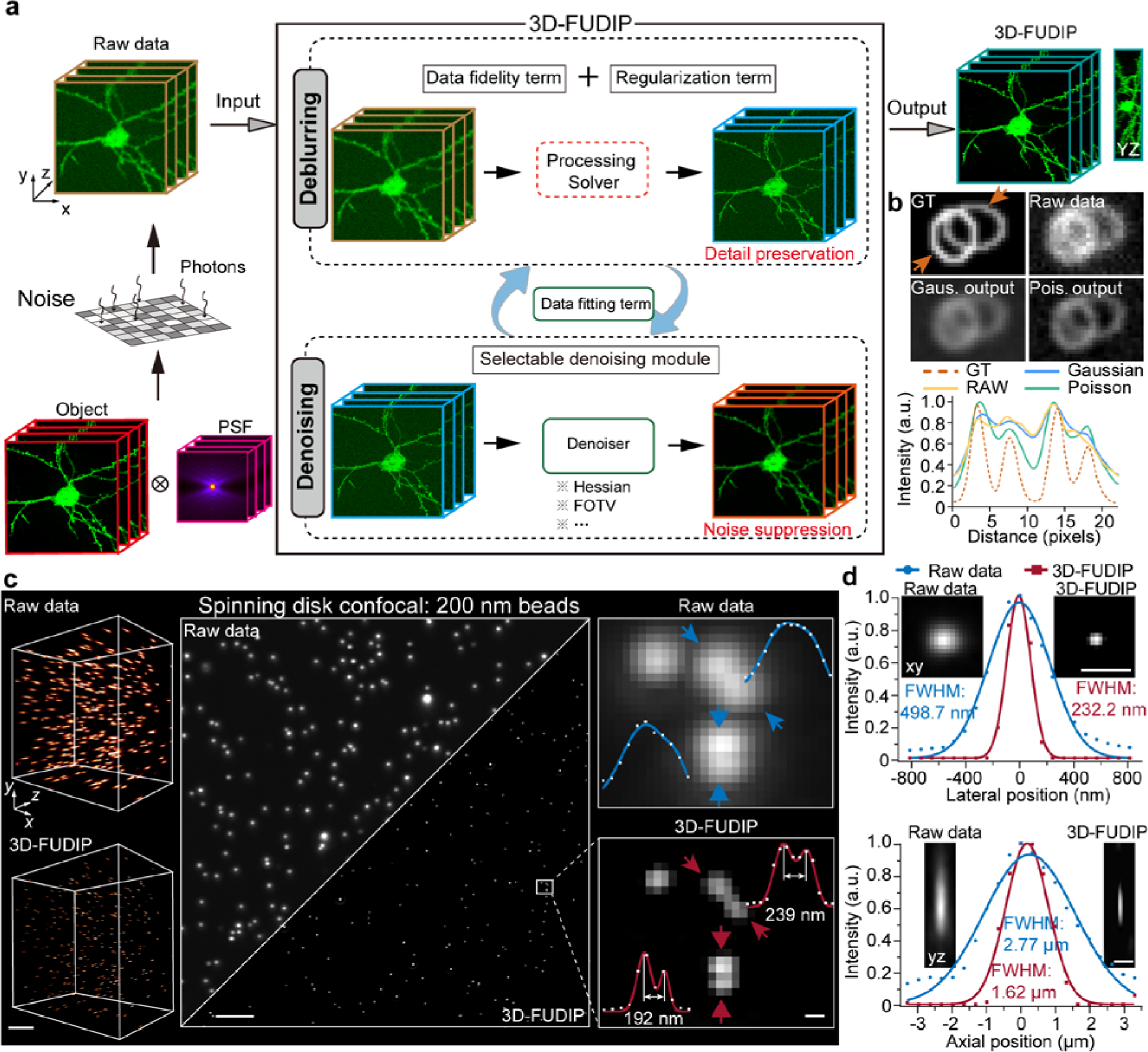
Principles of 3D-FUDIP. **a**, Workflow of 3D-FUDIP. 3D-FUDIP decouples the deconvolution into deblurring and denoising parts via variable splitting. The deblurring part can be solved through the alternating direction method of multipliers (ADMM) solver^36^. The denoising part is compatible with various denoisers. The results can be optimized with alternative iterations to achieve noise suppression and detail preservation, simultaneously. **b**, Results of the performance comparison between the Poisson model-based deconvolution algorithm and the Gaussian model-based deconvolution algorithm. **c**, The 3D volumes (left), maximum intensity projection (MIP) (middle), and enlarged ROIs (right) of 200-nm fluorescent beads. 3D-FUDIP resolved adjacent beads with distances of 192 nm, and 239 nm. **d**, The lateral FWHMs of raw data (blue) and 3D-FUDIP (red) are 498.7 nm and 232.2 nm, respectively. The axial FWHMs of the raw data (blue) and 3D-FUDIP (red) are 2.77 μm and 1.62 μm, respectively. Scale bars, 10 μm (**c** (left)), 50 μm (**c** (middle)), 200 nm (**c** (right)); 1 μm (**d**).

To investigate the imaging performance with 3D-FUDIP, we first imaged 200-nm fluorescent beads using a spinning disk confocal (SD-confocal) microscope with a 40× objective (UPLSAPO S, 40×/1.25-NA Silicone Oil, Olympus). Notable improvements have been achieved in spatial resolution and image contrast (Fig. 1c). The results show that 3D-FUDIP successfully resolved two beads with spacings of 192 nm and 239 nm in the lateral maximum intensity projection (MIP), achieving a near-diffracted-limited resolution (253.8 nm) of the microscope system equipped with a 40× objective. The lateral and axial full-width at half-maximum (FWHM) of the beads were decreased from 498.7 nm to 232.2 nm, and from 2.77 μm to 1.62 μm, respectively, achieving a nearly 8-fold improvement in spatial resolution (Fig. 1d).

### 3D-FUDIP enhances the SNR under low-light conditions

In many cases of biomedical imaging, there is severe noise interference especially in low light conditions, making it a challenge to restore sufficient details in the deconvolution process. 3D-FUDIP adopts the Poisson model, which obeys the quantum nature of light more than the conventional Gaussian-based model, making it suitable for imaging under different noise conditions, especially under low-light conditions. To evaluate the performance of 3D-FUDIP, we first generate 3D phantom objects^24^ consisting of spheres, ellipsoids, and cubes (Fig. 2a). Then the objects are convolved with the PSF of an SD-confocal microscope with a 40× objective. The final volumes contain both Poisson and Gaussian noises. The comparison results demonstrate that 3D-FUDIP outperforms conventional deconvolution algorithms with better noise suppression and detail restoration capability. The enlarged regions show that 3D-FUDIP can effectively identify the hollow rings that are indistinguishable in the noisy volume in both axial and lateral directions (Fig. 2b). The image quality enhancements can be quantified with SSIM and PSNR metrics (Fig. 2c). It can be noted that 3D-FUDIP outperforms conventional deconvolution algorithms at various noise levels, consisting with the above results. Even under the most severe noisy condition, 3D-FUDIP (SSIM=0.55, PSNR=24.71 dB) demonstrates substantial enhancement in comparison to the noisy blurred data (SSIM=0.13, PSNR=18.25 dB). In contrast, the results of Pure-let (SSIM=0.24, PSNR=18.96 dB), Huygens (SSIM=0.18, PSNR=19.03 dB), and RL (SSIM=0.20, PSNR=18.54 dB) only show minor improvements in SSIM and PSNR metrics.

**Fig. 2.**
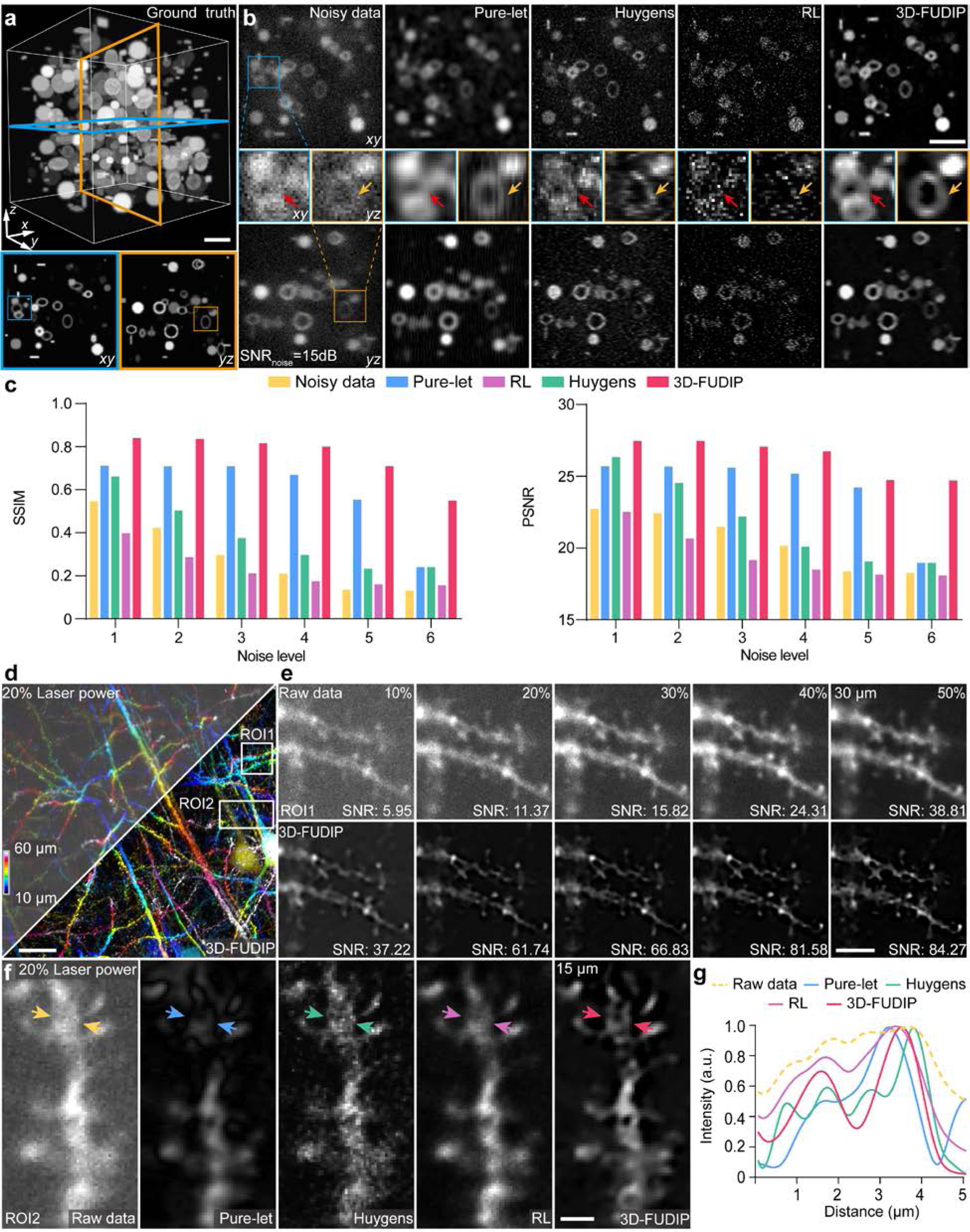
SNR enhancements with 3D-FUDIP under various noisy conditions. **a**, The 3D volume of the synthetic blurred phantom objects, which is regarded as the ground truth (GT) (noiseless and without blur). The lateral (left) and axial (right) cross-sections of the ground truth are shown at the bottom. **b**, The cross-section views of blurred noisy volume, Pure-let output, Huygens output, RL output, and 3D-FUDIP output are compared (left to right). The lateral cross-section (top row), enlarged ROIs (middle row), and axial cross-section (bottom row) are shown. **c**, Quantitative analysis for evaluating the performance of different algorithms under various noise conditions with SSIM and PSNR metrics. A larger value of noise level indicates a higher noise. **d**, Depth color-coded MIPs of raw data captured using 20% laser power (left) and the corresponding 3D-FUDIP output (right). **e**, Comparison between the raw data captured using different laser powers and the corresponding 3D-FUDIP outputs. From the left to right, we show the data captured by 10%, 20%, 30%, 40%, and 50% laser power, respectively. The enlarged ROIs with different laser powers correspond to the location of ROI1 in **d**. **f**, Comparison of results obtained using different deconvolution algorithms on data captured using 20% laser power. The enlarged ROIs correspond to the location of ROI2 in **d**. **g**, The intensity profiles between the arrows in **f**. Scale bars, 20 μm (**a**, **d**), 5 μm (**b**, **e**), 2 μm (**f**).

To further investigate the performance of 3D-FUDIP in biological samples, we imaged the Thy1-YFP mouse brain slices under various SNRs by adjusting the laser illumination power from 10% to 50% (Fig. 2d). The results show that 3D-FUDIP improves the SNR in various noisy conditions. Specifically, 3D-FUDIP achieves a 6-fold improvement (SNR_10%-3D-FUDIP_ = 37.22) at 10% excitation laser power, which is close to the SNR of raw data captured with 50% laser power (SNR_50%-power_ = 38.81). For high-quality images obtained with 50% excitation laser power, 3D-FUDIP still improves the SNR 2-fold (SNR_50%-3D-FUDIP_ = 84.27) (Fig. 2e). Compared with other deconvolution methods, 3D-FUDIP exhibits stronger noise robustness, enabling more fine structures to be clearly distinguished, especially in severe noisy conditions (Fig. 2f). The intensity profiles between the arrows across the dendrite confirm that 3D-FUDIP can recover a structure internal to the dendritic shaft, which is unresolved in raw data and other deconvolution results (Fig. 2g).

### 3D-FUDIP improves resolution and contrast with a switchable denoising module

3D-FUDIP has the advantage of fully decoupling the deconvolution process into two parts: deblurring and denoising. The deblurring part can be analytically solved through mathematical calculation, while the denoise part enables switchable denoisers to be embedded in the deconvolution process.

Further results captured by mouse brain slices verify that 3D-FUDIP is compatible with different denoisers (Fig. 3a, b). 3D-FUDIP successfully reversed the degradation caused by blur and noise and enhanced the contrast (Fig. 3c). Enlarged ROIs in different positions reveal that 3D-FUDIP resolves the hole structures and significantly extends the spatial frequency spectrum (Fig. 3d, e), with a nearly doubled lateral resolution quantified by the decorrelation method^37^ (Fig. 3f).

**Fig. 3.**
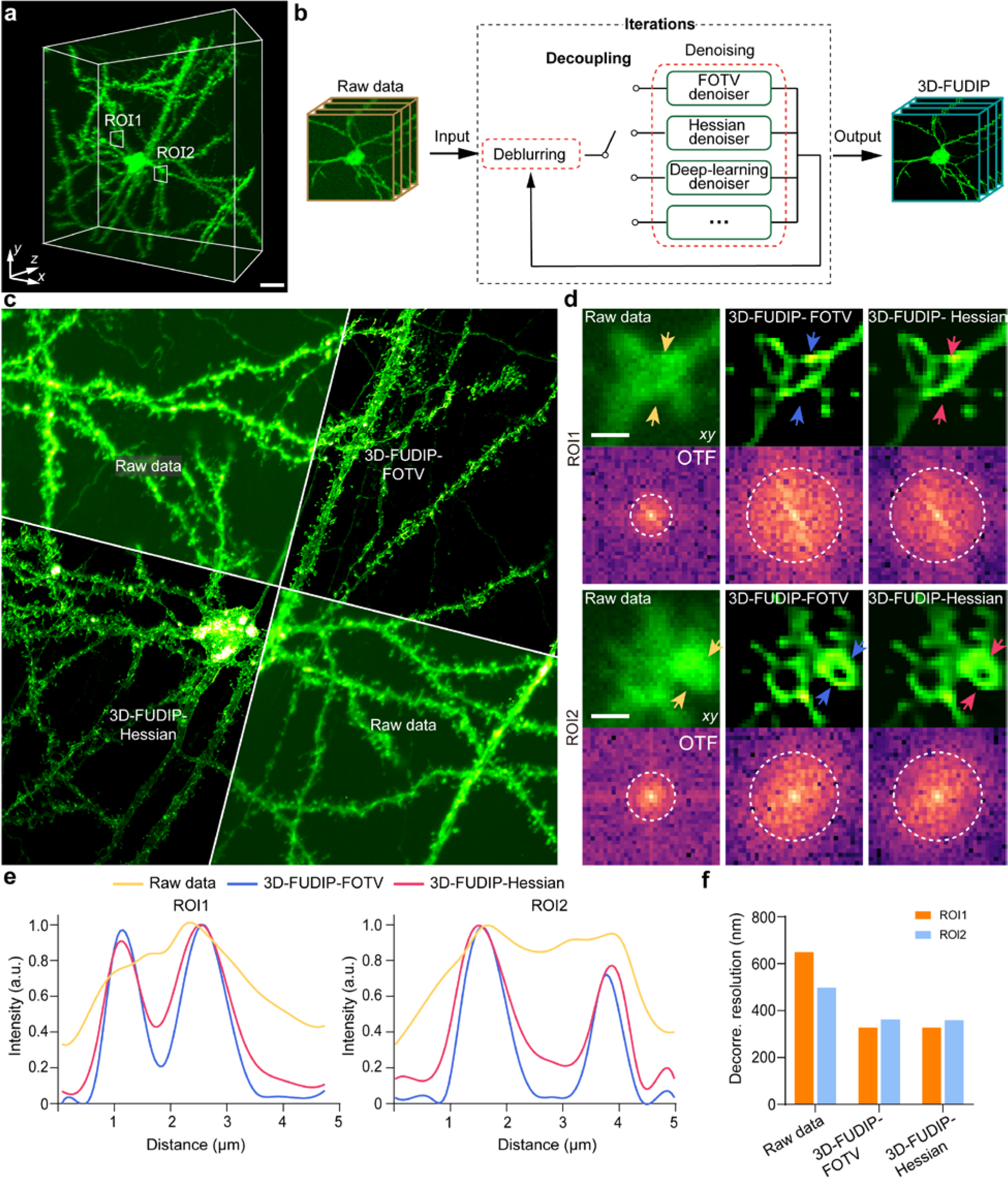
Resolution and contrast improvements by 3D-FUDIP with switchable denoiser. **a**, The primary somatosensory cortex (S1) mouse brain imaged by an SD-confocal microscope with a 20× objective. **b**, 3D-FUDIP with a switchable denoiser. **c**, Comparison of MIPs of raw data and 3D-FUDIP. **d**, Magnified ROIs and corresponding spectra. The locations of ROI1 and ROI2 correspond to the positions marked in **a**. **e**, The intensity profiles between the arrows in the ROI1 and ROI2. **f**, Decorrelation resolution analysis corresponds to ROI1 and ROI2. Scale bar, 20 μm (**a**), 10 μm (**c**), 1 μm (**d**).

### 3D-FUDIP boosts the space-bandwidth product

In biological imaging, the resolution and FOV are commonly a pair of contradictions^38, 39^. High-NA objectives offer superior resolution but have a small FOV and short working distance (WD). In contrast, low-NA objectives provide a larger FOV and longer WD at the expense of spatial resolution. The space-bandwidth product (SBP)^40^, which is defined as the product of image volume and bandwidth (1/resolution)^41^, is usually employed to quantify the comprehensive performance of resolution and FOV in an optical imaging system. Specifically, the maximum imaging volume of a 40× objective is almost 6 times larger than that of a 60× objective, while the resolution is worse (Fig. 4a). This trade-off between resolution and FOV should be considered when selecting an objective for a particular imaging application. Imaging the same area with different magnifications needs to be re-locate and refocus after replacing the immersion medium, leading to extra photobleaching and imaging efficiency reduction.

**Fig. 4.**
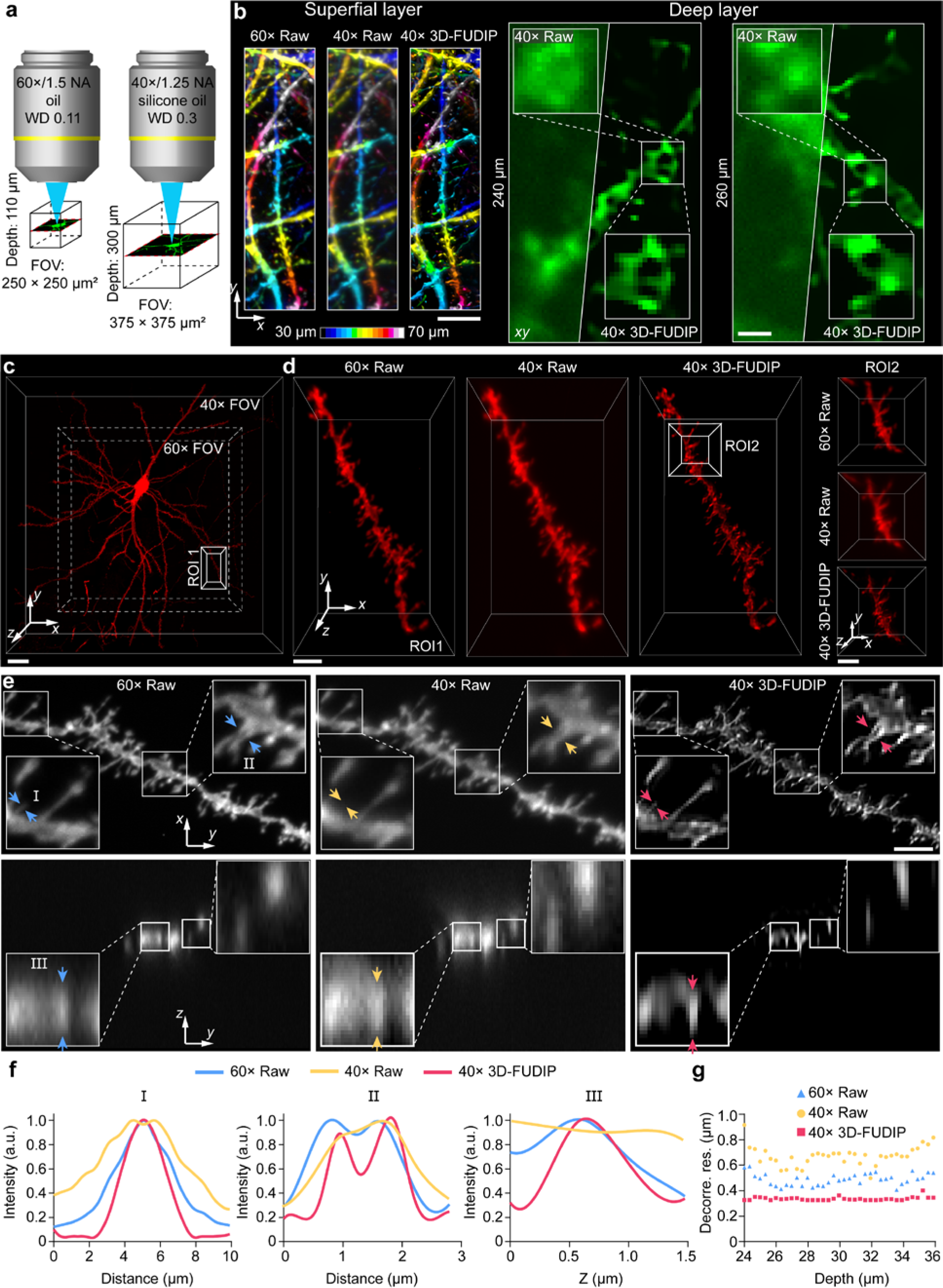
3D-FUDIP boosts the space-bandwidth product. **a**, Comparison of the working distance between the 40× objective and the 60× objective. **b**, Comparison of the depth color-coded MIPs among the 60× raw data (left), 40× raw data (middle), and 40× 3D-FUDIP (right) in the superficial layer of the mouse cerebral cortex. Comparison of lateral (*xy*) cross-sections between the 40× raw data and 40× 3D-FUDIP in the deep layer of the mouse cerebral cortex. **c**, The 3D volume of a sparsely labeled neuron was enhanced by 3D-FUDIP. **d**, ROI1: Magnified view of the white box in **c**. ROI2: Magnified views of the white box in **d**. **e**, The lateral (*xy*) MIPs of the corresponding dendrite segment in **d**. The lateral (*xy*) MIPs of 60× raw data (left in the top row), 40× raw data (middle in top row), and 40× 3D-FUDIP (right in top row) are shown in the top row. The ROIs inside the white rectangles are magnified. The axial (*yz*) views of 60× raw data (left in bottom row), 40× raw data (middle in bottom row), and 40× 3D-FUDIP (right in bottom row) are shown in the bottom row. **f**, The intensity profiles across the dendritic spine head in position I (left), different dendritic spines in position II (middle), and a dendritic spine along the *z* direction in position III (right). **g**, The decorrelation resolution at different axial positions in ROI1. Scale bars, 10 μm (**b** (left)), 2 μm (**b** (right)), 20 μm (**c**), 5 μm (**d**, **e**).

To solve this problem, we use 3D-FUDIP to boost the SBP by improving the resolution of images acquired with low-magnification objectives. By imaging 200-nm fluorescent beads, we show an increase in SBP from 30.61 million to 241.49 million by 3D-FUDIP with a 40× objective, which is roughly 20 times greater than that of a 60× objective (11.83 million). We then further applied 3D-FUDIP to images captured in the mouse cerebral cortex. The results demonstrate that the contrast and sharpness are significantly improved by 3D-FUDIP with a 40× objective, even outperforming those of a 60× objective acquired in the same area (Fig. 4b). Besides, when the imaging depth exceeds the WD of a 60× objective (110 μm), 3D-FUDIP can still realize good image restoration in deep layers (240 μm, 260 μm) with noticeable improvements (Fig. 4b).

To further investigate the improvements of SBP by 3D-FUDIP, we imaged neurons in mouse brain slice. (Fig. 4c, d). The results show substantial improvements in both lateral and axial resolution by 3D-FUDIP with a 40× objective, even surpassing those of a 60× objective (Fig. 4e). The unobserved dendritic spine neck and two adjacent dendritic spines are resolved after using 3D-FUDIP. Additionally, there are notable reductions in axial blur and background fluorescence (Fig. 4f). Finally, the resolution quantified by the decorrelation method^37^ at different axial positions shows that 3D-FUDIP with a 40× objective nearly doubles its resolution (from 673 nm to 337 nm), even surpassing that of a 60× objective (490 nm) (Fig. 4g).

### 3D-FUDIPn for 3D high-resolution imaging

To obtain further resolution enhancements, we proposed 3D-FUDIPn (Fig. 5a). Raw images were captured by an SD-confocal microscope with a 60× objective and further enhanced by 3D-FUDIP to generate high-resolution (HR) image volumes (regarded as GT). We can obtain degraded low-resolution (LR) images by imaging simulation.

**Fig. 5.**
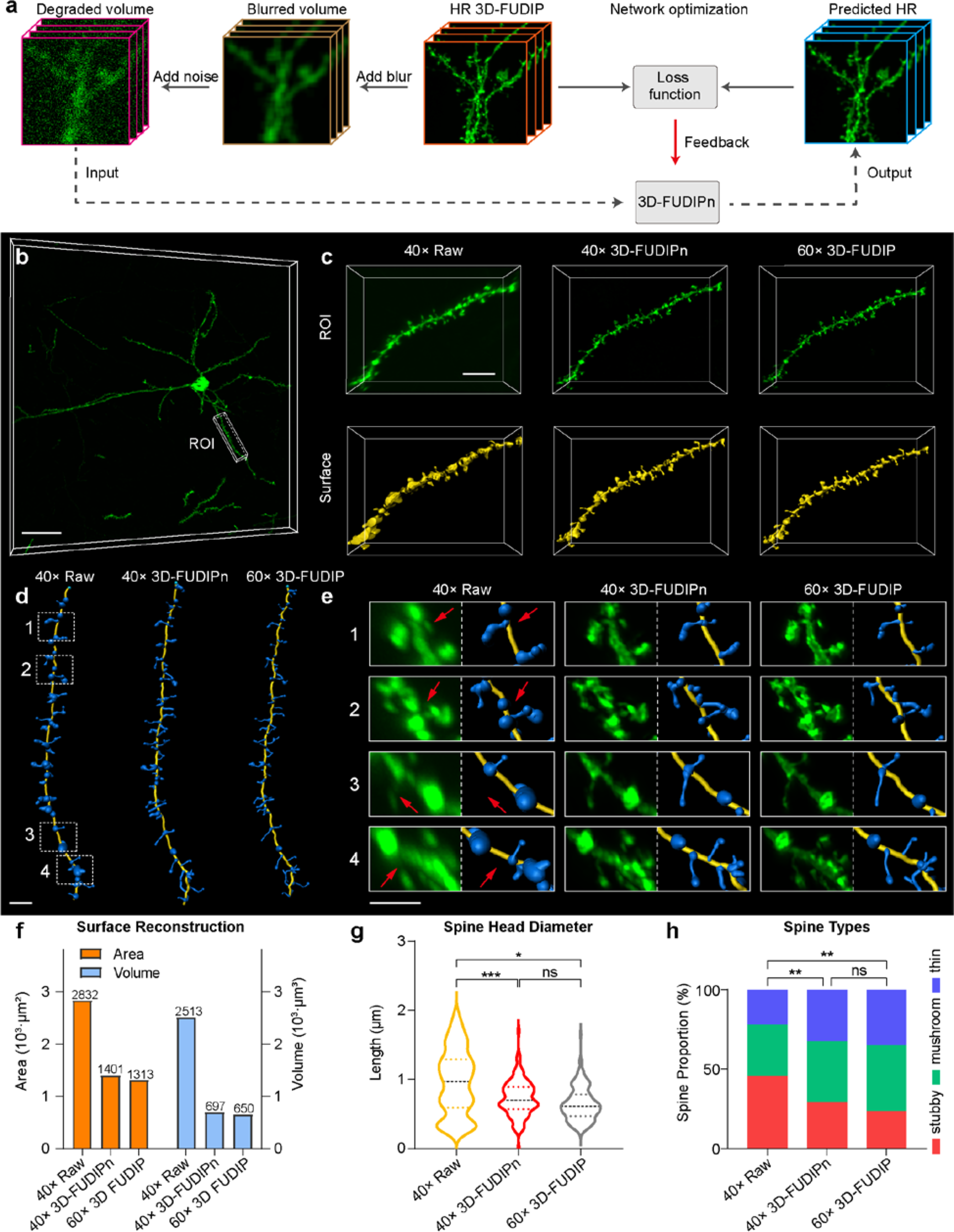
3D-FUDIP-based 3D-FUDIPn for 3D high-resolution imaging. **a**, The 3D high-resolution reconstruction by 3D-FUDIPn based on 3D-FUDIP. The high-resolution (HR) image volumes are achieved by 3D-FUDIP. The blurred and noisy low-resolution (LR) image volumes are synthetically generated by the degradation of HR. **b**, The imaging volume of a single neuron with a 60× objective. **c**, The ROIs and corresponding surfaces of 40× raw, 3D-FUDIPn with 40×, and 3D-FUDIP with 60× in the same area (left to right). **d**, Filament reconstruction of spines of 40× raw, 3D-FUDIPn with 40×, and 3D-FUDIP with 60× (left to right), corresponding to ROI in **b**. **e**, The enlarged ROIs of spines and corresponding filament reconstructions of 40× raw, 3D-FUDIPn with 40×, and 3D-FUDIP with 60× (left to right), corresponding to the white dashed rectangles in **c**. **f**, Statistical information of the dendrites in **b**. **g**, Statistical information of spine head diameters. (40× Raw:1.125±0.021 (mean ± SEM), spine number = 270; 3D-FUDIPn: 0.571±0.011 (mean ± SEM), spine number = 265; 60× 3D-FUDIP: 0.504±0.010 (mean ± SEM), spine number = 253; n=6 dendrites prepared from 3 mice.) **h**, Statistical information of spine types. (40× Raw: 54.4% stubby, 22.6% mushroom, 3.3% thin, 19.7% filopodia; 3D-FUDIPn: 21.0% stubby, 42.0% mushroom, 19.5% thin, 17.5% filopodia; 60× 3D-FUDIP: 15.2% stubby, 34.0% mushroom, 28.7% thin, 22.1% filopodia; n=270 spines prepared from 6 dendrites in 3 mice.) Scale bars: 50 μm (**b**), 20 μm (**c**), 5 μm (**d**), 10 μm (**e**).

To evaluate the performance of 3D-FUDIPn, we applied it to reconstruct the dendrites and spines in the primary somatosensory cortex (S1) (Fig. 5b, c). The enlarged ROIs of the reconstructed dendritic spines reveal that 3D-FUDIPn can clearly observe fine details that are unresolvable in the 40× raw data, thus enabling more accurate recognition of spines (Fig. 5d, e). The representative 3D volumes and automatically reconstructed spines prove that 3D-FUDIPn has good capability in reducing blur and suppressing background. The results of axial MIPs and intensity profiles indicate that the blurry spines are clearly resolved. The statistical results show that 3D-FUDIPn dramatically reduces the reconstruction error rates of the area and volume of dendrites by nearly 17-fold (40× Raw: 2832 μm^2^; 3D-FUDIPn: 1401 μm^2^; 60×3D-FUDIP: 1313 μm^2^) and 40-fold (40× Raw: 2513 μm^3^; 3D-FUDIPn: 697 μm^3^; 60× 3D-FUDIP: 650 μm^3^), respectively (Fig. 5f). Moreover, the spine head diameter distribution and mean values obtained by our method are more similar to the ground truth (Fig. 5g). Statistical information on spine type classification also presents a more accurate distribution by 3D-FUDIPn (40× Raw: 54.4% stubby, 22.6% mushroom, 3.3% thin, 19.7% filopodia; 3D-FUDIPn: 21.0% stubby, 42.0 % mushroom, 19.5% thin, 17.5% filopodia; 60×3D-FUDIP: 15.2% stubby, 34.0% mushroom, 28.7% thin, 22.1% filopodia).

## Discussion

The limited NA and noise inevitably degrade the imaging performance of optical microscopies. Deconvolution can partially enhance image details but is susceptible to noise, remaining a trade-off between detail preservation and noise suppression. Here, we develop 3D-FUDIP to break this trade-off by fully decoupling the deconvolution process to deblurring and denoising parts. By using 3D-FUDIP, we can improve the spatial resolution by nearly 8-fold and the spatial-bandwidth product (SBP) by 20-fold, allowing more details to be resolved within a larger imaging volume. By synergizing deep learning with these advances, we propose 3D-FUDIPn to further improve the imaging resolution. Besides, the adoption of the Poisson model enables more than a 6-fold enhancement in SNR with only a one-fifth photon budget, providing a potential solution for biomedical imaging with extremely low light conditions.

We demonstrate a doubled spatial spectrum, indicating enhanced detail recovery. The statistical results show that 3D-FUDIPn reduces the reconstruction error rates of dendrites by nearly 17-fold. Unlike expensive hardware-based imaging schemes, such as Mesolens^42^, RUSH^43^, and Schmidt objective^44^, 3D-FUDIPn conducts hardware-free image processing to achieve large FOV imaging with high resolution, providing a powerful tool for high-resolution computational imaging in biological science.

3D-FUDIP can also be applied to wide-field imaging systems such as LSFM. LSFM has the advantage of high imaging speed but suffers from relatively low resolution due to the low-NA objective and imperfect tissue clearing^45, 46^. By applying 3D-FUDIP to a home-built LSFM, we improve the detail preservation and spatial resolution (Fig. 6a, b). The dendrite tracing results demonstrate that longer dendritic branches and more dendritic terminals can be identified, which are overwhelmed by the background in the raw data (Fig. 6c, d).

**Fig. 6.**
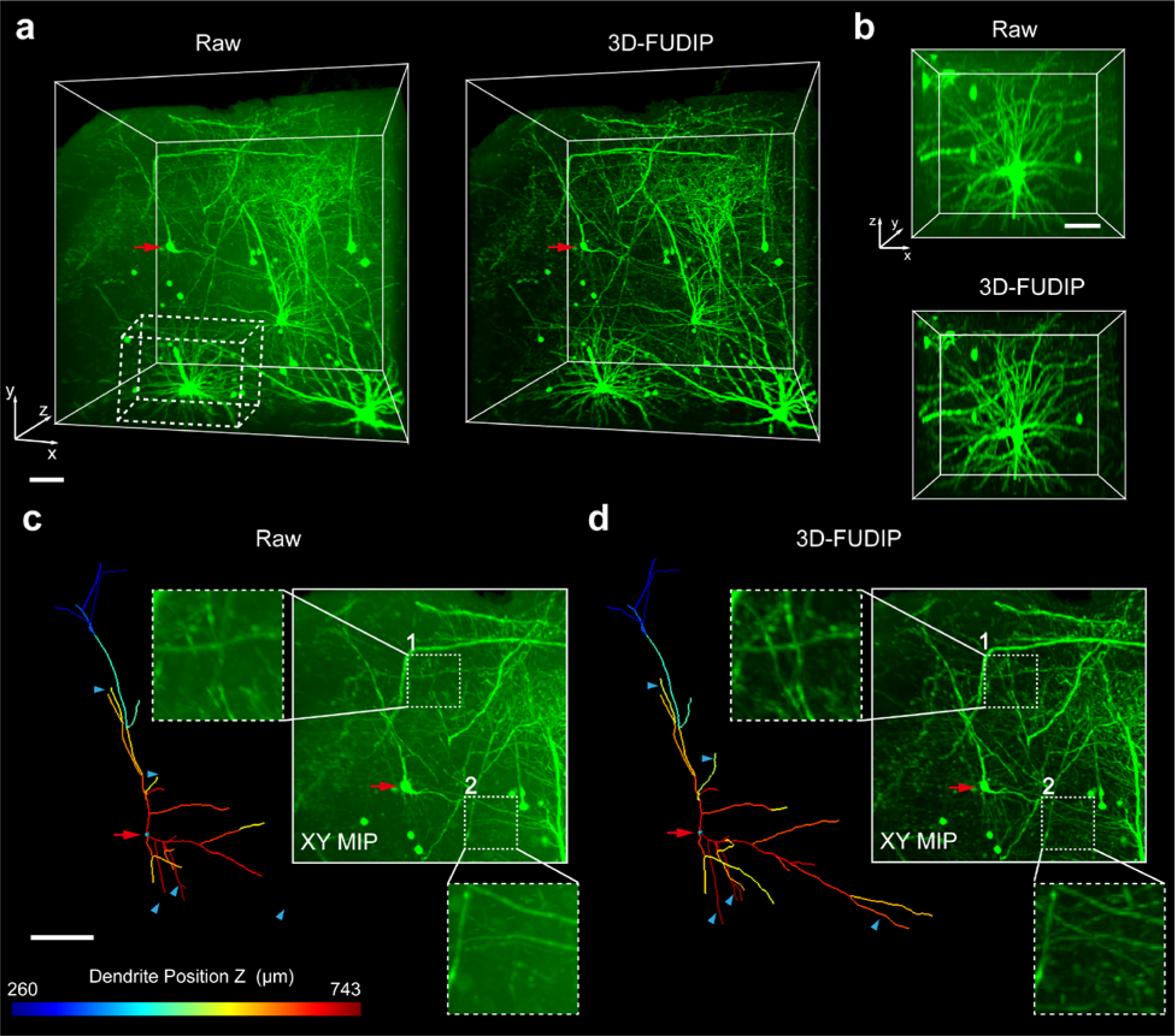
Compatibility of 3D-FUDIP with LSFM. **a**, A Thy1-YFP mouse brain tissue imaged by a home-built light-sheet microscope (left) and 3D-FUDIP (right). **b**, The enlarged ROIs of the raw data (top) and 3D-FUDIP (bottom), corresponding to the white dashed cube in **a**. **c**-**d,** The Imaris semi-automatic neuron tracing results and MIPs of raw data (**c**) and 3D-FUDIP (**d**). The red arrow highlights the location of the traced neuron soma. The blue triangles highlight the dendrite structures that are successfully reconstructed by 3D-FUDIP but are barely visible in the raw data. Scale bars: 100 μm (**a**-**d**).

We envision several extensions to our work. For the deblurring part, we currently assume a shift-invariant PSF approximation. In the future, we may extend our 3D-FUDIP to imaging systems with field-varying aberrations by employing spatially varying PSF. For the denoising part, we can embed networks with strong generalization and denoising capabilities into our framework, to further enhance the imaging performance. Besides, unsupervised learning might be integrated with our 3D-FUDIPn, reducing the reliance on high-quality datasets. With the development of computing power, our method also holds promise for applications with massive data, including large-scale volumetric imaging and long-term imaging, offering new insights not only for high-throughput 3D reconstruction but also live cell imaging with super-resolution under low-dose illumination.

## Code Availability

Custom codes for 3D-FUDIP computation developed in this study are available upon request from the corresponding author. The code will be released at https://github.com/WeLab-BioPhotonics/3D-FUDIP.

## Acknowledgements

This work was supported in part by STI 2030—Major Projects (2021ZD0200401), Key R&D Program of Zhejiang Province (2021C03001, 2022C03034), Natural Science Foundation of Zhejiang Province (LR22F050007), Non-profit Central Research Institute Fund of Chinese Academy of Medical Sciences (2023-PT310-01), Fundamental Research Funds for the Central Universities, Alibaba Cloud. We thank R. Fernandez for the help with 3D registration. We thank Shuangshuang Liu from the Core Facilities, Zhejiang University School of Medicine for their technical support. We thank Zizheng Wang and Yunyin Chen for sample preparation. We thank Tiancheng Zhang for assisting with video production.

## Contributions

K. S. and W. G. conceived the concept and supervised the research. Z.D. and J.C. developed the algorithms and programs. C.G. conducted the labeling experiments. C.G, Z.D. and J.C. performed imaging experiments. Z.D., C.G., J.C., and Y.Z. performed other experiments. Z.D. and J.L. developed the GUI. All the authors discussed the results and wrote the manuscript.

